# Hypoadiponectinemia does not enhance anxiety-like behaviour in a lean PCOS-like mouse model

**DOI:** 10.64898/2026.03.13.711682

**Authors:** Manisha Samad, Joakim Ek, Josefin Kataoka, Eva Lindgren, Claes Ohlsson, Ingrid Wernstedt Asterholm, Elisabet Stener-Victorin, Anna Benrick

## Abstract

Polycystic ovary syndrome (PCOS) is characterized by reproductive and metabolic disturbances and is associated with increased symptoms of anxiety and depression. Circulating adiponectin, an insulin-sensitizing adipokine, is reduced in women with PCOS, and low adiponectin has been linked to impaired mental health, particularly in females. We investigated whether low serum adiponectin is associated with impaired mental health in women with PCOS and whether adiponectin deficiency exacerbates anxiety-like behaviour in a PCOS-like mouse model.

Serum adiponectin was measured in women with (n=179) and without PCOS (n=228), stratified by body mass index (BMI). Health-related quality of life was assessed using the SF-36, generating physical and mental component scores. In parallel, the prenatal androgenization (PNA) PCOS-like mouse model was combined with adiponectin-deficient mice (APNhet) to assess the impact of reduced adiponectin on anxiety-like behaviour with and without prenatal androgen exposure.

Women with PCOS had lower total and high molecular weight adiponectin levels compared with controls. Adiponectin positively correlated with mental component scores in women with BMI <30, but not in those with obesity. Free testosterone was inversely correlated with adiponectin. In mice, PNA induced anxiety-like behaviour, however, reduced adiponectin did not exacerbate this phenotype. Although APNhet PNA mice showed 65% lower serum adiponectin levels and reproductive dysfunction, they displayed improved metabolic function. Unlike women with PCOS, adult PNA mice were not hyperandrogenic.

These findings suggest that adiponectin is associated with mental health in non-obese women, but reduced adiponectin alone does not induce anxiety-like behaviour in the absence of hyperandrogenism. The differing patterns observed across BMI categories, as well as between the human cohort and experimental data, underscore the complexity of the mechanisms underlying mental health disturbances in PCOS.

## Introduction

Polycystic ovary syndrome (PCOS) is a common endocrine disorder affecting approximately 11-13% of women of reproductive age [1]. It is a multifaceted condition with reproductive, metabolic, as well as psychological manifestations [1]. The predominant characteristics include hyperandrogenism, ovulatory dysfunction and polycystic ovarian morphology. Clinically, PCOS is often associated with hirsutism, menstrual irregularities, decreased fertility, obesity and increased risk of type 2 diabetes [1]. According to the current diagnostic criteria, PCOS is defined by the presence of at least two of the following three features: 1) clinical/biochemical hyperandrogenism, 2) ovulatory dysfunction, and 3) polycystic ovaries on ultrasound or elevated anti-mullerian hormone (AMH) levels [1].

Beyond its reproductive and metabolic consequences, PCOS is strongly associated with impaired metal health [2]. Screening for anxiety and depression is therefore recommended at the time of diagnosis [3]. However, many patients report that psychological issues are under-recognized, and report dissatisfaction with the support and counselling received [4]. While much of the research on PCOS has focused on its reproductive and metabolic features, women with PCOS also report significantly lower mental health–related quality of life [2, 5]. They also experience a high burden of psychological symptoms, with studies reporting depressive symptoms in approximately 30–40% of women and anxiety symptoms in roughly 30–75%, depending on the population studied and the assessment method used [6–8]. This could be due to the social stigma of PCOS-symptoms such as obesity, infertility, acne, and hirsutism, along with the chronic, complex, and frustrating nature of PCOS, contributing to depression and anxiety symptoms [9]. In addition, anxiety and depressive symptoms have been linked to hyperandrogenism and insulin resistance, suggesting underlying biological contributors beyond psychosocial stressors [10]. Although PCOS is a heritable disorder, currently identified genetic variants explain only approximately 10% of the estimated 70% heritability, indicating that PCOS develops due to a combination of genetic and environmental factors [11]. Moreover, summary statistics from genome-wide association studies suggest no shared genetic basis or causal relationship of PCOS with psychiatric disorders including depression, anxiety, schizophrenia and bipolar disorder, suggesting that the association is likely mediated by BMI [12].

Both clinical and preclinical studies highlight the role of excess androgens in the development of PCOS [2, 13, 14]. In pregnant women with PCOS, elevated androgen levels persist throughout gestation and androgens can cross the placenta, potentially influencing fetal development and the development of PCOS symptoms [15–17]. Prenatal androgen exposure has been shown in both human and animal studies to affect cognitive and behavioural outcomes in female offspring [17, 18]. We and others have demonstrated that prenatal androgen exposure leads to the development of anxiety-like behavior in female rodents [13, 17]. Consistent with these findings, children of women with PCOS are at increased risk of neuropsychiatric disorders, including anxiety, depression, ADHD, and autism spectrum disorder [18]. Using Swedish population-based health registers, we have reported an 78% increased risk of anxiety diagnosis among daughters of women with PCOS [17]. Together, these findings suggest that maternal androgen excess may contribute to long-term neuropsychiatric vulnerability in female offspring.

Another potential mediator linking PCOS and impaired mental health is reduced adiponectin levels. Adiponectin is an adipocyte-produced insulin-sensitizing hormone with important metabolic properties [19]. Emerging evidence suggests sex-specific interactions between severe mental disorders and circulating levels of testosterone, sex hormone–binding globulin (SHBG), and adiponectin, with women exhibiting higher testosterone and lower adiponectin levels in association with psychiatric conditions, pointing to a potential sex-dependent pathophysiology [20]. In women with PCOS, adiponectin levels are lower and correlates with insulin resistance [21, 22]. A meta-analysis further demonstrated an inverse association between circulating adiponectin and psychiatric disorders, including anxiety, mood, and trauma-related disorders [23]. Notably, the association with depressive disorders was stronger in females [20]. Together, these findings suggest that adiponectin may represent a biological link between metabolic dysfunction, hyperandrogenism, and mental health vulnerability in women.

However, the mechanisms through which adiponectin and androgens influence mental health remain poorly understood. We therefore investigated whether low serum adiponectin is associated with impaired mental health symptoms in women with and without PCOS and if reduced adiponectin exacerbates anxiety-like behaviour in a PCOS-like mouse model. To mechanistically examine the interaction between adiponectin and anxiety, PNA model was combined with adiponectin-deficient mice. The PNA model develops an anxiety-like behaviour and is characterized by PCOS-like reproductive and metabolic disturbances [17, 24].

## Material and methods

### Clinical cohort

Data from four studies conducted in Sweden have been used for this study [25–28] resulting in a total 407 women aged 18-38 years (PCOS n= 179, non-PCOS n= 228). A summary of each study is described in **Supplemental table 1**. The studies included women in a wide range of BMIs, recruited from the community or an obesity unit between 2005-2016. The patients were screened for PCOS with the Rotterdam-criteria (two out of three criteria; hyperandrogenism, irregular cycles, and/or polycystic ovary morphology) except for the severely obese cohort [28] where the National Institute of Health (NIH) criteria (hyperandogenism and irregular cycles) was used, as ultrasound was not a successful method to quantify polycystic ovary morphology. Ethical approval for the four studies was received from the Ethics Committee, University of Gothenburg, Gothenburg, Sweden. All studies were registered at clinical-trials.gov; NCT01319162, NCT00484705, NCT01457209, and NCT00921492. The included studies were performed in accordance with the Declaration of Helsinki, and all participants gave their oral and written informed consent to participate in the study.

### Data collection

Data on symptoms of anxiety and depression and health-related quality of life (HRQoL) were assessed from the questionnaires Comprehensive Psychopathological Rating Scale (CPRS) Self-rating Scale for Affective Syndromes (CPRS-S-A), and short form (SF)-36. To assess symptoms of anxiety and depression we used the CPRS-S-A. It is a validated and well-established instrument used both in research and clinical routine psychiatric care for defining symptoms of anxiety and depression and for evaluation of treatment. From CPRS-S-A, the subscales Brief Scale for Anxiety (BSA) to assess symptoms of anxiety, and Montgomery Åsberg Depression Rating Scale (MADRS-S) to assess symptoms of depression, are extracted [29, 30]. The scales consist of nine domains each, of which two are present in both scales. All domains are rated on a six-point scale: 0 represents an absence of symptoms; 2 represents a potentially pathological deviation; 4 represents a pathological condition and 6, an extremely pathological condition. Domain ratings are summarized, and a maximum value of 54 for each scale is calculated. A cut-off value of >11 in the BSA-S and the MADRAS-S scales was used to set a reference value for clinically relevant anxiety and depression.

HRQoL is a way of describing personal health status, including physical and mental health. To assess HRQoL, SF-36 questionnaire was used. It is generic and designed to capture a person’s perception of how his or her health status has interfered with physical, psychological, and social functioning for the past four weeks, and is a validated instrument widely used [31]. It is divided into eight domains: physical functioning, physical role limitation, bodily pain, general health perception, emotional role limitation, vitality, mental health, and social functioning. In each domain a score of 0-100 is calculated, where 0 represents the worst possible health quality and 100 represents the best possible health quality. Summary scores were computed to reflect the physical component score (PCS) and mental component summary (MCS), scoring on a scale of 0-100.

BMI was calculated using weight in kilos, divided with squared height in meters. Hirsutism was assessed with Ferriman-Gallwey (FG)-score. Serum samples were collected in the morning from subjects fasted overnight. Total and high molecular weight (HMW) adiponectin levels were measured by ELISA (80-ADPHU-E01, Alpco, Salem, NH, USA). Serum samples were thawed, diluted and treated according to the manufacturer’s instructions. The absorbance was measured using a plate reader (SpectraMax iD3, Molecular Devices, San Jose, CA, USA). Testosterone, SHGB, and insulin were analyzed by ECLIA (CBAS 8000 Roche Diagnostics Scandinavia AB, Sweden) at the ISO-accredited laboratory at Sahlgrenska University Hospital, Gothenburg, Sweden. Free testosterone was calculated using total testosterone and SHBG according to Vermeulen et al [32]. Free androgen index (FAI) was calculated as (total testosterone (nmol/L) / SHBG (nmol/L) x 100.

### Animal Model

A mouse model with decreased adiponectin levels was used to study the effect of adiponectin, with and without PNA. Wt and APNhet mice on a C57BL/6 background were used [33]. All the animals had ad libitum access to food and water and were maintained under standard housing conditions (12-hour light/dark cycle environment, and fixed temperature and humidity) provided by the Laboratory of Experimental Biomedicine, University of Gothenburg. The study was approved by the Ethics Committee of the University of Gothenburg (ethical number) in accordance with the ARRIVE guidelines, the European Union guidelines for the care and use of laboratory animals (2010/63/EU), and the Swedish Board of Agriculturés regulations and general advice of laboratory animals (L150).

### Experimental design

APNhet males were mated with wt females and gave birth to both wt and APNhet offspring allowing us to study littermates (see breeding scheme and study outline in **Fig.1**). Before mating, vaginal smears were performed to determine the estrous cyclicity of the female mice [34]. The females in estrous phase were allowed to mate with APNhet males overnight. The weight was monitored to confirm for pregnancy associated weight gain. At GD 16.5 to 18.5, vehicle or 250μg 5α-Androstan-17β-ol-3-one (dihydrotestosterone (DHT), A8380, Sigma-Aldrich, St. Louis, USA) dissolved in 2.5μl benzyl benzoate (B6630, Sigma-Aldrich) and 47.5μl sesame oil (S3547, Sigma-Aldrich) was injected subcutaneously in the interscapular area. The breeding resulted in 53 mice divided into 4 groups: wt or APNhet offspring from vehicle injected wt dams (wt veh, n=9, APNhet veh, n=10) or DHT-injections during gestation to induced PNA (wt PNA, n=22, APNhet PNA, n=10). Female offspring were weaned at 3 weeks of age and genotyped as previously described [35]. Ear biopsies were incubated with DirectPCR lysis buffer (Viagen #102-T) and Proteinase K solution (Invitrogen Direct PCR #25530-049, 0.2 mg/mL) at 55^0^C overnight and then at 80^0^C for 45 minutes to inactivate the enzyme. Lysate was centrifuged at 14,000 × *g* for 3 min. Two microliters of each sample was mixed with 18 µL master mix containing PCR Supermix (ThermoFisher, #10572014) with the following primers: Wt-forward, 5′-TTCAATTCCAG-CACCCACAGTAA-3′; Wt-reverse, 5′-GGACCCCTGAACTTGCTTCAC-3′; and APNhet-forward 5′-GTAGCCGGATCAAGCGTATG-3′; and APNhet-reverse, 5′ATGAACTCCAGGACGAGGCA-3′. The PCR was performed with an enzyme activation step, 5 min at 95 °C, then 30 sec at 95 °C, 30 s at 58 °C, 1 min at 72 °C for 35 cycles. The resulting PCR products were run separately on a 2% agarose gel, and the mice were genotyped by the presence of a 350-bp amplicon for the APNhet allele and a 350-bp amplicon for the wild type allele.

**Figure 1:**
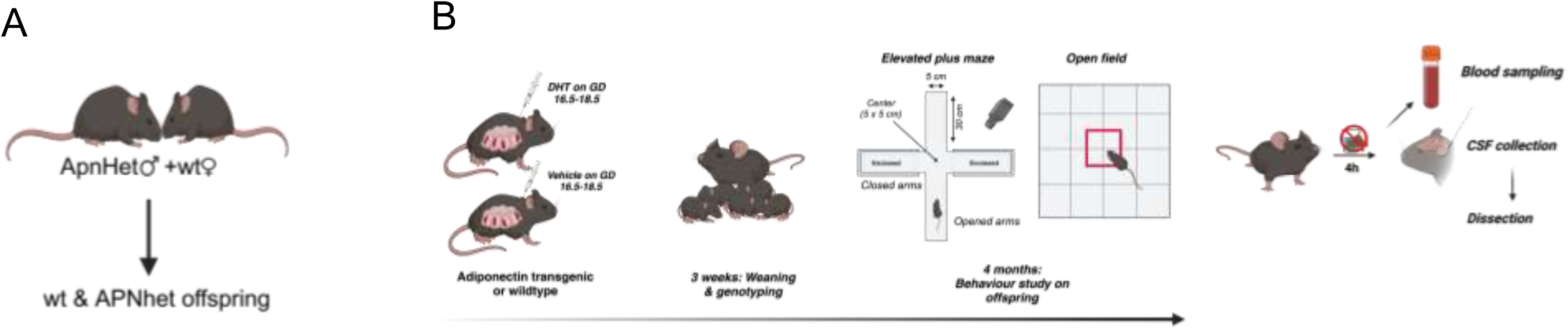
Study design. **(A)** Schematic representation of the breeding scheme generating wildtype (wt) & adiponectin heterozygous (APNhet) littermates. **(B)** describes the overall study plan where the effects of adiponectin deficient with and without prenatal androgenization (PNA) on anxiety-like behavior were studied using elevated plus maze and open field followed by dissection. Dihydrotestosterone; DHT (created with https://BioRender.com).

### Behavioural tests

Behavioural tests were carried out at 4 months of age. Before starting the experiment, all mice were allowed to acclimate for one hour in the room where the experiments were conducted. Anxiety-like behaviour was evaluated using an elevated plus maze (EPM) comprised of two open and two closed arms (45x10cm) and a platform in the middle (10x10cm), elevated 50 cm above the ground. The mouse was placed in the junction and allowed to move freely for five minutes, which was recorded using a camera mounted above the maze. The maze was placed into a dimly lit room so that light intensity reached 10-15 lux in the closed arms and 50-55 lux in the open arms. Movement was immediately tracked and prepared for further analysis using EthoVision 13 XT. Data were evaluated for time spent in the closed and open arms of the maze, time spent in the middle, number of entries in the arms, and distance travelled. This test is based on the innate aversion of mice against bright and open spaces and mice are more anxious when they spend less time in the open arms of the apparatus.

As an additional measurement of anxiety-like behaviour, mice were placed in a brightly lit open filed (OF, 100x100x40cm) and allowed to freely explore for 15 minutes. Levels of anxiety were assessed by locomotor activity and the amount of time spent in the central part of the arena. The OF was lit with a light intensity of 35 lux. Movement was recorded with a camera mounted above the arena and tracked with EthoVision XT, data were evaluated for time spent in the central area of the OF and distance travelled. This test, like EPM, is also based on the innate aversion of mice against bright and open spaces. The animal is considered more anxious when it spends less time in the open area of the apparatus.

### Tissue Collection

The behavioural tests were followed by 5 days of vaginal smears to check estrous cyclicity. The mice were fasted for 3-4 h before a small tail blood sample was taken to quantify serum insulin levels by ELISA (Mouse insulin 10-1247-01, Mercodia, Sweden) and glucose was measured with a blood glucose meter (Contour XT, Bayer Health Care, Mishawaka, USA). Mice were then sedated with isoflurane followed by heart puncture, and a larger blood sample for serum analyses was collected along with inguinal adipose tissue (IWAT), gonadal adipose tissue (GWAT), and ovaries. In a subset of mice, cerebrospinal fluid (CSF) was collected by puncturing the cisterna magna using a glass capillary with a fine tip and gentle suction [36]. Samples with suspected blood contamination were immediately discarded. Total adiponectin was analyzed in serum (diluted 1:50 000) and CSF (diluted 1:100) using an adiponectin ELISA (80569, CrystalChem, Zaandam, Netherlands) with a sensitivity of 0.008ng/mL, with an intra-assay coefficient variation (CV)< 10%.

AMH was measured in the serum using AMH ELISA kit (AL-115, AnshLabs). The absorbance was measured on a plate reader (SpectraMax I3x, Molecular Devices, San Jose, CA, USA). Serum concentrations of sex hormones were determined with a validated liquid chromatography/mass spectroscopy (GC–MS/MS) [37].Values below lowest level of quantification (LLOQ) were excluded from summary calculations.

### Ovary histology

Ovaries were collected and immediately fixed in 4% paraformaldehyde for 24-48 hours. They were transferred to 70% ethanol and dehydrated with different concentrations of ethanol and embedded in paraffin. Later, the ovaries were sectioned at a thickness of 5 𝜇m in a microtome. Tissues were deparaffinized with xylene and then rehydrated in graded ethanol baths (100%, 90%, 80%, 70% and 50%) followed by rinses in distilled water. Finally, they were stained with haematoxylin & eosin. The number of preantral and antral follicles, and corpus luteum were counted under a microscope [34].

### Statistical analyses

SPSS 27.0 was used for statistical analyses. Normality of data was assessed by Kolmogorov-Smirnov statistics. The distribution of scores was not normally distributed in the clinical cohort; therefore Mann-Whitney U-test was used to compare women with and without PCOS. Adjustment for age was calculated with ANCOVA. Correlation analyses between adiponectin, mental health scores, and free testosterone were made using Spearman’s rank correlation test.

The effect of PNA on behavior, ovary histology and AMH in wt mice was analysed by an unpaired Student’s t-test. Moreover, wt PNA mice were compared to APNhet PNA to determine if there was an additive effect of adiponectin deficiency. Differences in metabolic characteristics, serum and CSF adiponectin levels compared to wt veh was analyzed using a one-way analysis of variance (One-way ANOVA). Data are expressed as mean ± SEM and *P < 0.05* was considered significant.

## Results

### Clinical cohort

Detailed anthropometrics data and mental health scores from the four studies have been previously published [25–28, 38]. Normal weight and overweight women with PCOS in this study have previously been shown to have more symptoms of depression and lower MCS compared with controls, and normal-weight women with PCOS have more symptoms of anxiety [38]. Based on this finding, we divided the cohort into BMI<30 kg/m^2^ and BMI≥30 kg/m^2^. Secondary analyses of PCOS-characteristic variables and mental health scores divided into BMI<30 kg/m^2^ and BMI≥30 kg/m^2^ are presented in **Table 1**. Women with PCOS have higher FG-scores, total testosterone, free testosterone and FAI, and lower SHBG. Weight, BMI, and fasting insulin did not differ between groups (**Table 1**). In those with BMI<30 30 kg/m^2^, symptoms of depression and anxiety was higher and MCS was lower in women with PCOS compared to controls (**Table 1**).

**Table 1:** Metabolic and psychological characteristics of women with PCOS and controls stratified by BMI (<30 and ≥30 kg/m^2^).

| Variable | BMI <30<br>Controls<br>(n=37) | BMI <30<br>PCOS<br>(n=76) | P-value | BMI ≥30<br>Controls<br>(n=191) | BMI ≥30<br>PCOS<br>(n=103) | P-value | Adj.<br>P-value |
| --- | --- | --- | --- | --- | --- | --- | --- |
| Age (y) | 27.7 | 28.8 | 0.194 | 37.6 | 32.5 | <0.001 |  |
| Weight (kg) | 69.8 | 67.1 | 0.287 | 110.3 | 105.7 | 0.015 | 0.059 |
| BMI (kg/m<sup>2</sup>) | 24.9 | 23.8 | 0.132 | 39.4 | 38.2 | 0.032 | 0.081 |
| FG score | 0.4 | 10.6 | <0.001 | 3.2 | 12.3 | <0.001 | <0.001 |
| SHBG (nmol/L) | 67.3 | 47.0 | <0.001 | 33.2 | 27.2 | <0.001 | <0.001 |
| Total testosterone (nmol/L) | 0.78 | 1.34 | <0.001 | 1.11 | 1.62 | <0.001 | <0.001 |
| Free testosterone (nmol/L) | 0.0096 | 0.0213 | <0.001 | 0.0209 | 0.0331 | <0.001 | <0.001 |
| FAI | 1.47 | 3.73 | <0.001 | 3.95 | 6.97 | <0.001 | <0.001 |
| F-insulin (mU/L) | 6.68 | 7.32 | 0.194 | 17.61 | 19.19 | 0.028 | 0.102 |
| CPRS-SA Anxiety | 9.19 | 14.40 | <0.001 | 16.60 | 16.00 | 0.689 | 0.786 |
| CPRS-SA Depression | 6.76 | 13.20 | <0.001 | 13.60 | 13.40 | 0.891 | 0.983 |
| SF-36 MCS | 46.3 | 32.3 | <0.001 | 38.1 | 36.6 | 0.335 | 0.551 |

Data are shown for controls and women with PCOS within each BMI category. Comparisons between controls and women with PCOS were performed using Mann–Whitney *U* test. Adjusted *P* values were obtained by ANCOVA with adjustment for age. Abbreviations: PCOS, polycystic ovary syndrome; FG score, Ferriman–Gallwey score; SHBG, sex hormone-binding globulin; FAI, free androgen index; f-insulin, fasting insulin; CPRS-SA, Comprehensive Psychopathological Rating Scale–Self-Assessment; SP-36 MCS, 36-Item Short Form Survey Mental Component Summary.

Total and HMW adiponectin were lower in women with PCOS compared to controls (**Figure 2A**), with no difference in the HMW/total adiponectin ratio (0.332 ± 0.011 vs 0.338 ± 0.011, *P* = 0.51).

**Figure 2:**
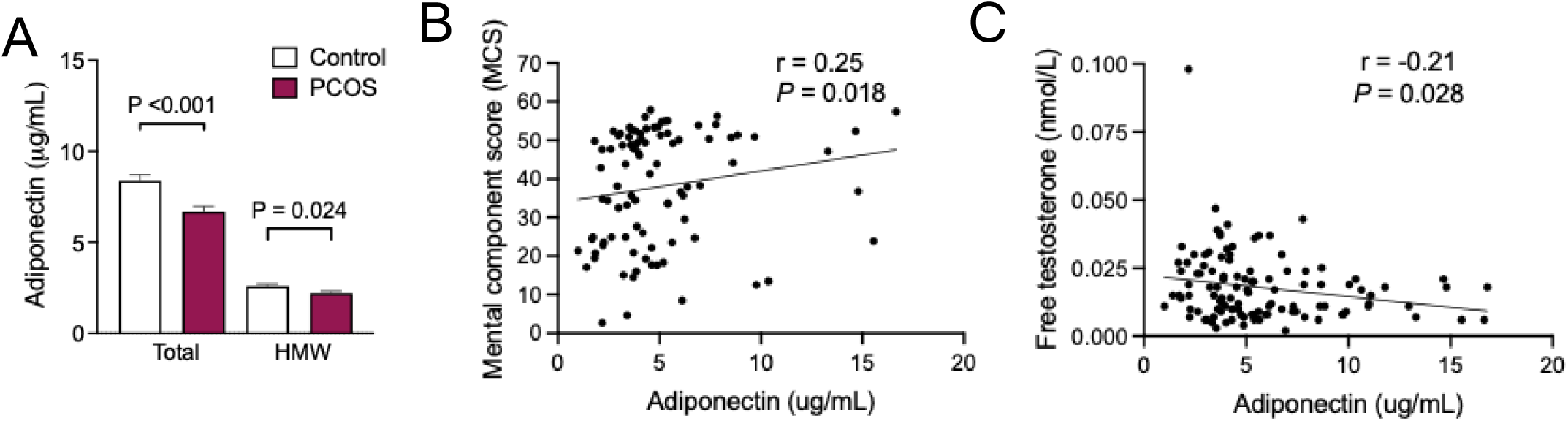
Clinical cohort. **(A)** Total and HMW adiponectin in women with PCOS (n= 179) and controls (n= 228), **(B)** Correlation between serum adiponectin and mental component summary score and (**C**) between serum adiponectin and free testosterone in women with BMI<30 (n=113). Mann-Whitney U-test was used to compare women with and without PCOS. Correlations were analyzed with Spearman’s rank correlation test. Data presented as mean ± SEM.

There were no differences in symptoms of anxiety and depression, or MSC in women with obesity and severe obesity [38], therefore we performed correlation analyses including only women with BMI<30 kg/m^2^ to determine if adiponectin correlates with mental health scores. Higher total adiponectin correlated with MCS (**Figure 2B)**, while there was no correlation between adiponectin and anxiety (-0.052, *P*=0.58) or depression (-0.090, *P*=0.34). Testosterone inhibits adiponectin expression and secretion from adipocytes *in vitro* [39, 40]. In line with this, increased free testosterone (**Fig. 2C)** and free androgen index (FAI) correlated with lower adiponectin *(r*= -0.20, *P* = 0.037).

### Behaviour characteristics

An adiponectin deficient mouse model was used to take the above finding from correlation to causality. Adiponectin deficient mice were used to determine if lower adiponectin enhances the anxiety like-behaviour in prenatal androgenized mice.

APNhet, being an adiponectin deficient model, showed a 65% decrease in serum compared to wt **(Figure 3A)**. A similar effect was seen in CSF, confirming that central adiponectin was also decreased in APNhet mice **(Figure 3B)**. These was a strong correlation between serum and CSF levels *(r = 0.65, P<0.0001)*.

**Figure 3:**
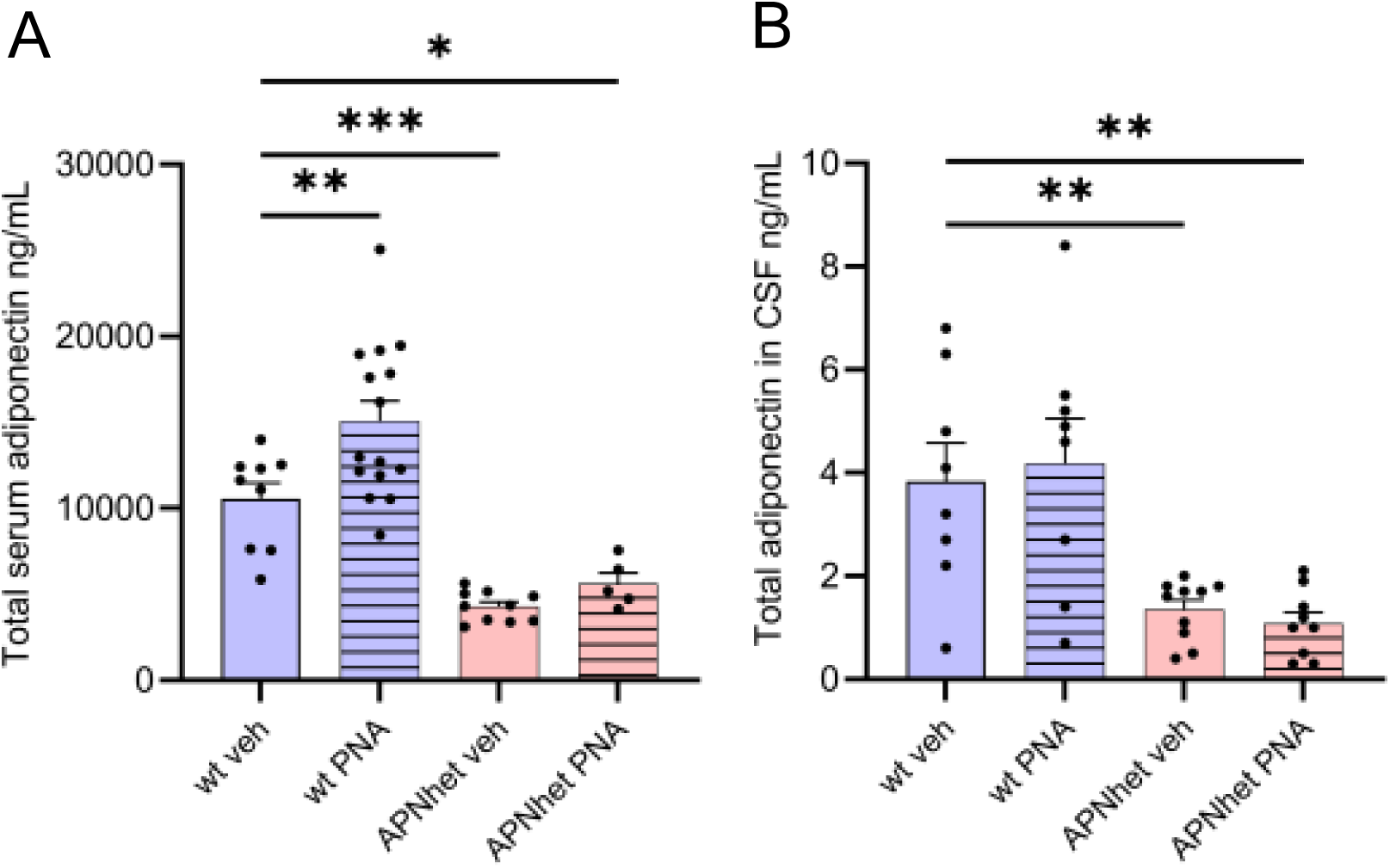
Serum measurements. **(A)** Total serum adiponectin in 4-month-old wild-type (wt) and adiponectin heterozygous (APNhet) female mice with and without prenatal androgenization (PNA), (**B)** total adiponectin in CSF measured in wild-type (wt) and adiponectin heterozygous (APNhet) female mice. Differences in serum and CSF adiponectin compared to wt veh was analyzed using One-way ANOVA. Data are expressed as mean ± SEM and *\*P<0.05, **P<0.01,* \*\*\**P < 0.001*

As expected, wt PNA mice spent less time in the open arms and tended to spend more time in the closed arms, indicating anxiety-like behaviour compared to the wt veh group (**Figure 4A, B**). APNhet PNA mice spent more time in the centre compared to wt PNA **(Figure 4 C)**. The number of entries in the closed arms were also lower in wt PNA offsprings, this would indicate that the animals preferred to stay in the closed arms **(Figure 4D).** APNhet mice exhibit normal locomotor activity and there was no evident difference in behaviour in unchallenged or PNA mice compared to wt, except for number of entries in the closed arms which were lower in APNhet PNA **(Figure 4A-D)**. There was no difference in the time spent in the centre zone, or the number of entries into the centre zone, or the distance moved in the open field between groups (**Supplement table 2**).

**Figure 4:**
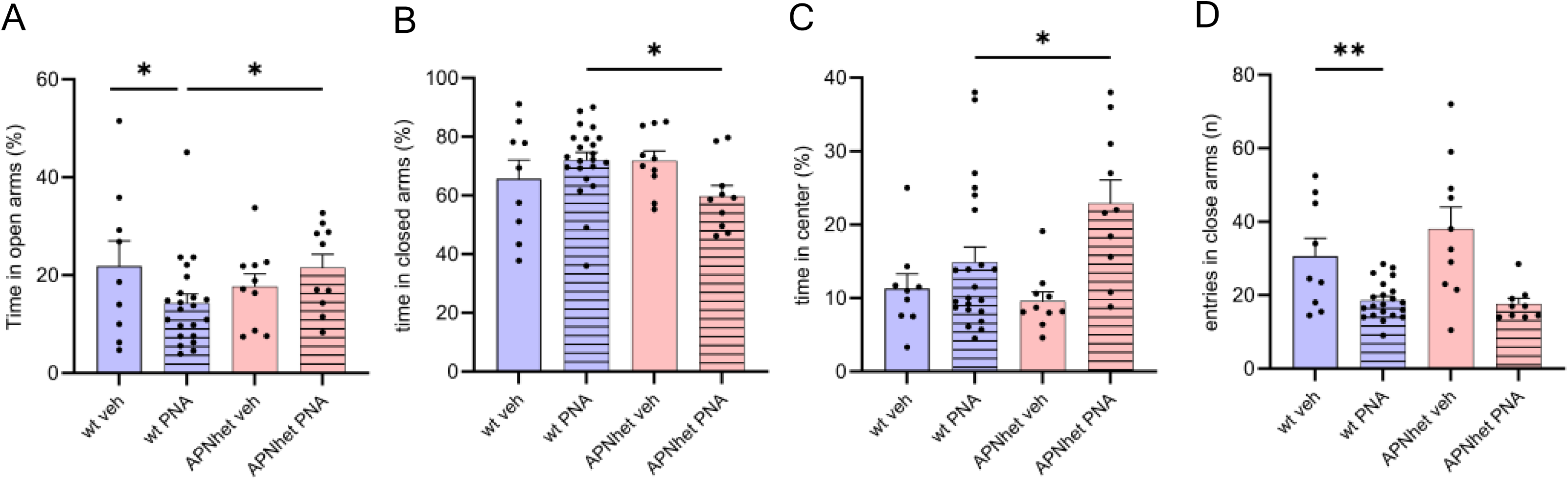
Behaviour characteristics EPM. **(A)** Time spent in open arms, **(B)** Time spent in closed arms, **(C)** Time spent in centre, and **(D)** number of entries into closed arms in 4-month-old wild-type (wt) and adiponectin heterozygous (APNhet) female mice with and without prenatal androgenization (PNA). The effect of PNA in wt mice was analysed by an unpaired Student’s t-test. wt PNA mice were compared to APNhet PNA to determine if there was an additive effect of adiponectin deficiency. Data are expressed as mean ± SEM and *\*P < 0.05, **P < 0.01*

### Metabolic and reproductive characteristics

There was no difference in body, IWAT and GWAT weights between the groups **(Table 2)**. Fasting glucose levels were reduced in the APNhet groups compared with the wt veh group, while fasting insulin levels tended to be lower in the PNA groups, although this difference was not statistically significant. (wt PNA *P*=0.054, APNhet PNA *P=0.052)* **(Table 2)**.

**Table 2:**
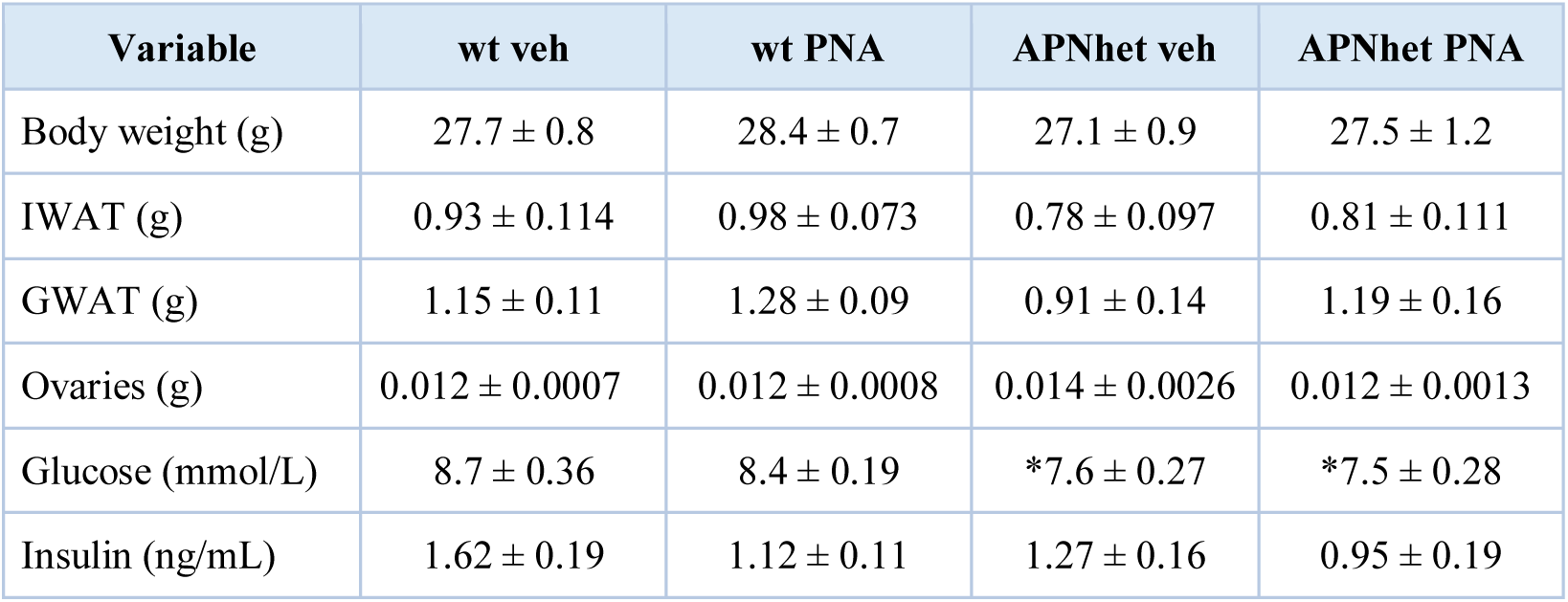
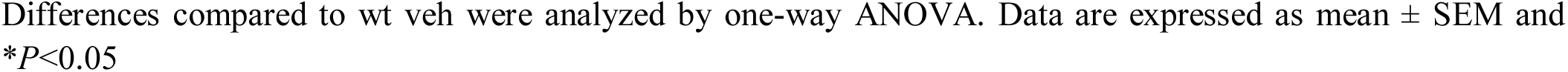
Metabolic characteristics of 4-month-old wild-type (wt) and adiponectin heterozygous (APNhet) female mice with and without prenatal androgenization (PNA).

| Variable | wt veh | wt PNA | APNhet veh | APNhet PNA |
| --- | --- | --- | --- | --- |
| Body weight (g) | 27.7 ± 0.8 | 28.4 ± 0.7 | 27.1 ± 0.9 | 27.5 ± 1.2 |
| IWAT (g) | 0.93 ± 0.114 | 0.98 ± 0.073 | 0.78 ± 0.097 | 0.81 ± 0.111 |
| GWAT (g) | 1.15 ± 0.11 | 1.28 ± 0.09 | 0.91 ± 0.14 | 1.19 ± 0.16 |
| Ovaries (g) | 0.012 ± 0.0007 | 0.012 ± 0.0008 | 0.014 ± 0.0026 | 0.012 ± 0.0013 |
| Glucose (mmol/L) | 8.7 ± 0.36 | 8.4 ± 0.19 | *7.6 ± 0.27 | *7.5 ± 0.28 |
| Insulin (ng/mL) | 1.62 ± 0.19 | 1.12 ± 0.11 | 1.27 ± 0.16 | 0.95 ± 0.19 |

Differences compared to wt veh were analyzed by one-way ANOVA. Data are expressed as mean ± SEM and \* $P < 0.05$

PNA wt mice had an increased number of preantral follicles and a trend towards more antral follicles **(Figure 5A).** An increase in antral follicles is often associated with polycystic ovarian morphology and anovulation in women with PCOS. However, no differences were seen in ovary weight or the number of corpora lutea **(Figure 5A**, **Table 2)**. We measured AMH in the serum as a marker for polycystic ovary morphology. In wt mice, PNA exposure tended to increase AMH concentrations relative to wt veh; however, this difference did not reach statistical significance *(P = 0.071)* **(Figure 5B)**. Sex hormone levels did not differ between groups (**Table 3**).

**Figure 5:**
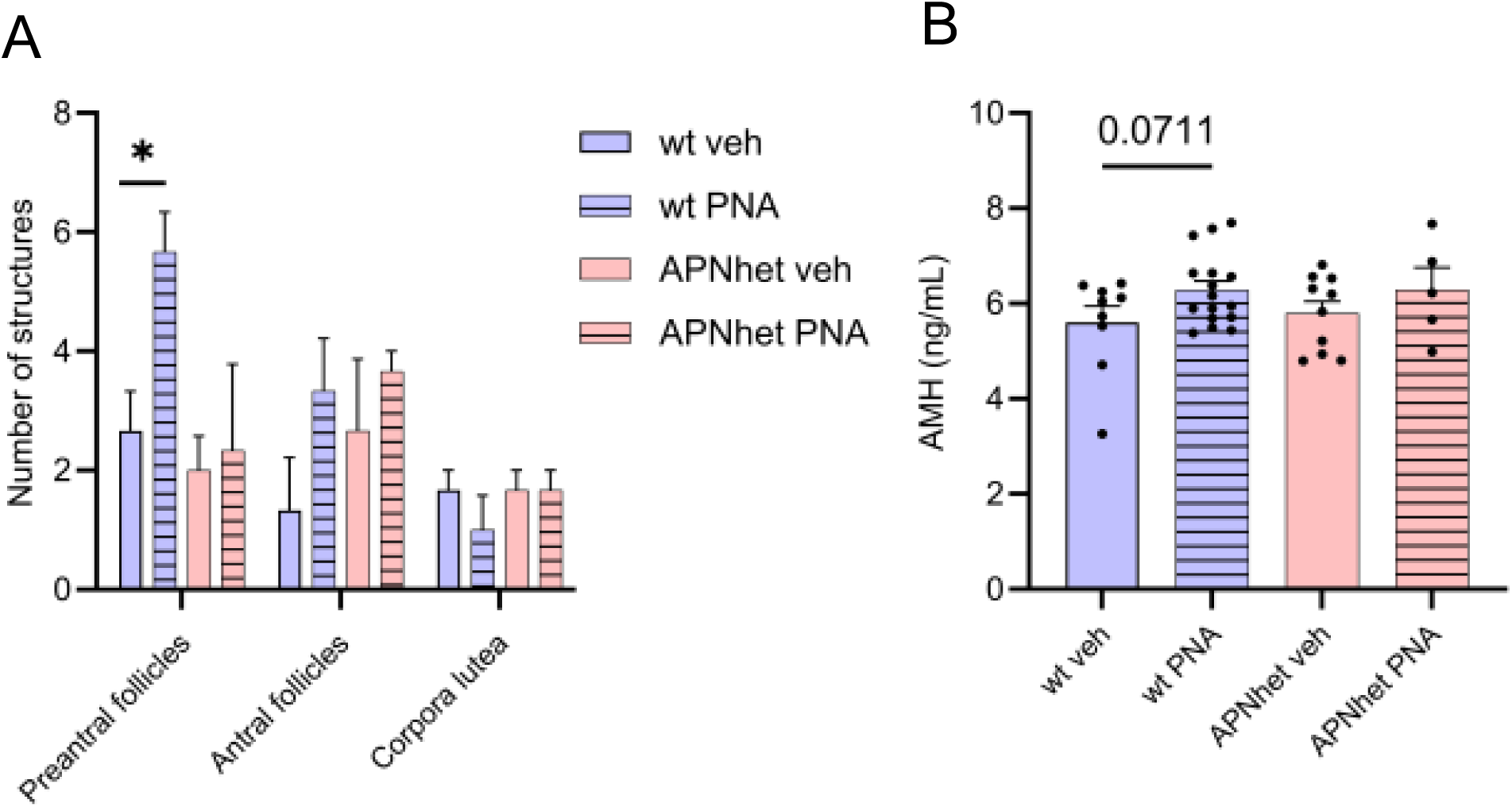
Ovary histology. **(A)** Quantitative analyses of the mean number of preantral, antral & corpora lutea (CL), n=4/group **(B)** Serum Anti-mullerian hormone (AMH) levels in 4-month-old wild-type (wt) and adiponectin heterozygous (APNhet) offsprings with and without prenatal androgenization (PNA). The effect of PNA in wt mice was analysed by an unpaired Student’s t-test. wt PNA mice were compared to APNhet PNA to determine if there was an additive effect of adiponectin deficiency. Data are expressed as mean ± SEM and \**P < 0.05*

**Table 3:** Serum concentrations of sex hormones in 4-month-old wild-type (wt) and adiponectin heterozygous (APNhet) female mice with and without prenatal androgenization (PNA).

| Variable | wt veh | wt PNA | APNhet veh | APNhet PNA |
| --- | --- | --- | --- | --- |
| Androstenedione (pg/mL) | 48.2 ± 4.4<br>n = 10 | 46.1 ± 4.2<br>n = 10 | 50.0 ± 10.3<br>n = 10 | 51.9 ± 4.4<br>n = 10 |
| Progesterone (pg/mL) | 5351 ± 1356<br>n = 10 | 2627 ± 673<br>n = 10 | 4808 ± 1779<br>n = 10 | 3676 ± 2267<br>n = 10 |
| DHT (pg/mL) | 9.7 ± 1.7<br>n = 3 | –<br>n = 0 | 7.1 ± 2.6<br>n = 6 | 6.5 ± 0.0<br>n = 1 |
| Estradiol (pg/mL) | 3.18 ± 1.10<br>n = 9 | 1.68 ± 0.46<br>n = 8 | 3.78 ± 1.22<br>n = 10 | 3.04 ± 0.74<br>n = 9 |
| Testosterone (pg/mL) | 17.2 ± 3.7<br>n = 8 | 11.3 ± 1.2<br>n = 3 | 21.9 ± 5.5<br>n = 6 | 27.5 ± 5.8<br>n = 8 |

Hormone concentrations measured by gas chromatography–tandem mass spectrometry (GC–MS/MS). Data are presented as mean ± SEM. Differences compared to wt veh were analyzed by one-way ANOVA. n=10/group. If n<10, excluded samples were below the LLOQ.

Serum testosterone and DHT were below the LLOQ in many mice, especially in the PNA groups, thus many missing values. This contrasts with the clinical cohort where women with PCOS are hyperandrogenic.

## Discussion

Women with PCOS report higher levels of anxiety and depressive symptoms and HRQoL compared with women without PCOS [8, 41, 42]. In the present study, lower serum adiponectin was associated with poorer mental health scores, suggesting that reduced adiponectin may contribute to the increased psychological burden observed in PCOS. We also found an inverse correlation between adiponectin and free testosterone, consistent with previous evidence that androgens suppress adiponectin secretion from adipocytes [39, 40]. Although both testosterone and adiponectin are well known for their peripheral metabolic effects, they can also cross the blood–brain barrier and are detectable in cerebrospinal fluid, indicating potential central actions [43, 44]. To disentangle the mechanistic relationship between CSF adiponectin and mental health outcomes, we used a genetically modified mouse model with low adiponectin levels [33]. Contrary to our hypothesis, reduced adiponectin did not exert an additive effect on androgen-induced anxiety-like behaviour. These findings suggest that while adiponectin is associated with mental health measures in women with PCOS, it may not independently potentiate androgen-driven anxiety-related behaviour, at least within the experimental framework used here.

Symptoms of depression and anxiety arise from complex interactions between brain circuits and neurotransmitter systems [45]. For testosterone and adiponectin to influence such behaviors, they likely need to access and act within the central nervous system. According to the Developmental Origins of Health and Disease (DOHaD) hypothesis, conditions during fetal life can program long-term disease risk in adulthood [46]. Daughters of women with PCOS are exposed *in utero* to higher androgen levels [15], which may constitute an initial “hit” affecting brain development and rewiring of brain circuits. As these daughters later develop PCOS themselves [47, 48], persistent hyperandrogenism and reduced adiponectin during reproductive life may represent a second exposure, potentially reinforcing these early changes [49]. The PNA mouse model does not fully replicate this endocrine milieu. Wt dams were used for breeding, exposing fetuses to elevated androgens but normal maternal adiponectin. As maternal adiponectin does not cross the placenta [50], and fetal adiponectin production in mice begins late in gestation (around embryonic day 17) at relatively low levels [51], prenatal programming in this model is likely driven primarily by androgen exposure. This differs from humans, where fetal adiponectin production begins in the second trimester, coinciding with adipose tissue development [52].

In adulthood, PNA APNhet offspring display reduced adiponectin but unaltered circulating androgens. Although adiponectin levels were threefold lower in the cerebrospinal fluid of APNhet mice, this reduction did not exacerbate PNA-induced anxiety-like behaviour. While the adipose tissue–brain axis has emerged as a potential modulator of anxiety, and adiponectin has been suggested to exert anxiolytic effects [53, 54], our findings do not support a direct anxiogenic effect of reduced adiponectin in this model. Moreover, the PNA model has imprinting effect on somatic tissues, including the brain, but it does not lead to hyperandrogenism in adult mice. Species differences in fetal adiponectin dynamics, as well as the absence of adult hyperandrogenism in the PNA model, may contribute to the discrepancy between the human cohort and the experimental data. The lack of a more pronounced anxiety-like phenotype in adiponectin-deficient mice may therefore reflect the requirement for concurrent hyperandrogenism to unmask such effects.

Higher insulin levels have been associated with increased symptoms of anxiety and depression in women, both with and without PCOS [38]. Studies have shown that insulin resistance in the brain can disrupt dopamine signaling and contribute to mood disorders [55]. As insulin resistance and impaired glucose homeostasis can influence brain function through altered insulin signaling in the CNS, low-grade inflammation, and changes in neurotransmitter systems [56, 57], we used a lean mouse model to minimize the cofounding effect of metabolic dysfunction. The PNA model did not exhibit insulin resistance, on the contrary, wt PNA mice showed unaltered glucose and tended to have lower fasting insulin levels. which contrasts with previous studies reporting androgen-induced glucose intolerance [58]. Notably, the PNA APNhet group displayed both lower fasting glucose and insulin levels, indicating improved glucose homeostasis. This unexpected finding may be explained by increased Uncoupling protein-1 (UCP-1) expression in brown and a trend towards increased UCP-1 in white adipose tissue as this activation has been linked to enhanced insulin sensitivity and glucose utilization in mice [59, 60]. Consequently, the relatively favorable metabolic profile of APNhet mice may have attenuated neurobiological changes typically associated with metabolic dysfunction, thereby masking potential anxiogenic effects of hypoadiponectinemia.

Higher BMI has been associated with increased symptoms of anxiety and depression in women with PCOS [61]. In our cohort, however, this association was observed only among women with BMI <30. In women with obesity, anxiety and depression scores were comparable between those with and without PCOS [38], suggesting that obesity may override the additional impact of hyperandrogenism on mental health. Consistent with this, the positive correlation between adiponectin and mental component scores was present only in women with BMI <30. In the highest BMI group, adiponectin levels did not differ between women with and without PCOS, further supporting the notion that obesity per se exerts a stronger influence on metabolic and psychological health than PCOS status alone. Clinically, these findings suggest that BMI should be carefully considered when evaluating mental health in women with PCOS. In women with BMI<30, hyperandrogenism and adipokine dysregulation may play a more prominent role in mood symptoms. Future interventional studies should examine whether metabolically focused treatments can alleviate mood symptoms and whether treatment responses differ according to BMI and endocrine phenotype.

In conclusion, serum adiponectin is associated with mental component scores in women, suggesting a link between adipokine regulation and psychological well-being. However, reduced adiponectin alone, in the absence of hyperandrogenism and hyperinsulinemia, did not induce anxiety-like behaviour in our mouse model. The differing patterns observed across BMI categories, as well as between the human cohort and experimental data, underscore the complexity of the mechanisms underlying mental health disturbances in PCOS. Together, our findings point to a multifactorial interplay between adiponectin, testosterone, and obesity in shaping mental health outcomes. Further studies are needed to clarify how these factors interact across developmental stages and metabolic contexts to influence vulnerability to anxiety and depression in PCOS.

## Acknowledgements

We thank the Experimental Biomedicine (EBM) core facility for their help with animal housing maintenance and Professor Karolina P Skibicka for the behaviour setup at Sahlgrenska Academy, University of Gothenburg.

## Author contributions

AB and MS conceived and designed the experiments. AB, MS, JE, JK, and EL performed the laboratory work and analyzed the data. CO performed the sex-hormone measurements and analysis. AB, IWA, and ESV supervised the project. AB and IWA secured funding. AB and MS took the lead in writing the manuscript. All authors provided critical feedback and contributed to the final approval of the manuscript.

## Data Availability

The data supporting the findings of this study are available on request with the corresponding author.

