## Supplement table 1 and 2 for "Hypoadiponectinemia does not enhance anxiety-like behaviour in a lean PCOS-like mouse model"

**Supplement Table 1**: Characteristics of studies included in analyses.

| Study | 1) Kataoka et al.[22] | 2) Jedel et al.[20] | 3) Benrick et al.[19] | 4) Johansson et al.[21] |
|---|---|---|---|---|
| Year | 2011-2016 | 2005-2008 | 2011-2013 | 2009-2010 |
| Registered at clinical trials.gov | NCT01319162 | NCT00484705 | NCT01457209 | NCT00921492 |
| Study design | Case-control study | Case-control study | Case-control study | Baseline measures of a randomized controlled trial |
| Number (PCOS/non-PCOS) | 246 (63/183) | 60 (30/30) | 42 (21/21) | 32 (32/0) |
| Age | 18-50 | 18-37 | 18-38 | 18-38 |
| BMI | PCOS: 39.9 ±4.7<br>Non-PCOS: 39.6 ±4.3 | PCOS: 24.8 (18.2-40.3)<br>Non-PCOS: 24.7 (19.3-41.6) | PCOS: 31.6 ±4.12<br>Non-PCOS: 30.41 ±3.62 | PCOS: 24.7 ±3.1<br>Non-PCOS: 23.2 ±3.6 |
| Location | Sweden | Sweden | Sweden | Sweden |
| Recruitment | Obesity unit, Sahlgrenska university hospital | Community | Community | Community |
| Diagnostic criteria | NIH<br>(>35 days cycle or <8 menstruations/year and clinical (FG>6) or biochemical hyperandrogenism (T >2.1nmol/L or fT >0.035 nmol/L or FAI >5) | Rotterdam<br>2/3 criteria of:<br>Oligo-/amenorré (35 days cycle or <6 menstruations/year or amenorrhea)<br>Polycystic ovaries (>12 follicles of 2-9 mm and/or ovarian volume of > 10 ml in one or both ovaries<br>Clinical hyperandrogenism (acne or hirsutism or FG-score>8) | Rotterdam<br>2/3 criteria of:<br>Oligo-/amenorré (35 days cycle or <6 menstruations/year or amenorrhea)<br>Polycystic ovaries (>12 follicles of 2-9 mm and/or ovarian volume of > 10 ml in one or both ovaries<br>Clinical hyperandrogenism (acne or hirsutism or FG-score>8) | Rotterdam<br>2/3 criteria of:<br>Oligo-/amenorré (35 days cycle or <6 menstruations/year or amenorrhea)<br>Polycystic ovaries (>12 follicles of 2-9 mm and/or ovarian volume of > 10 ml in one or both ovaries<br>Clinical hyperandrogenism (acne or hirsutism or FG-score>8) |
| Exclusion criteria | Pregnancy, breastfeeding last six months, language barrier. | Any pharmacological treatment last 12 wk, breastfeeding last 24 wk, self-reported physical or psychiatric disease, language barrier | Pharmacological treatment last 3 months, acupuncture last 2 months, breastfeeding last 6 months, DM type 1, cardiovascular disease, daily smoking/alcohol consumption | BMI<30, pharmacological treatments last 3 months, breastfeeding/acupuncture last 24 wk, cardiovascular disease, DM, language barrier |

**Supplemental Table 2:** Open field behavior measurements in 4-month-old wild-type (wt) and adiponectin heterozygous (APNhet) female mice with and without prenatal androgenization (PNA).

| | **wt veh** | **wt PNA** | **APNhet veh** | **APNhet PNA** |
|---|---|---|---|---|
| Distance moved (cm) | 7801 ± 850 | 9901 ± 664 | 8309 ± 617 | 8985 ± 892 |
| Time in center (%) | 5.43 ± 0.83 | 5.41 ± 0.63 | 5.95 ± 0.86 | 8.20 ± 1.49 |
| Entries in center (n) | 23.1 ± 3.3 | 31.5 ± 3.6 | 31.1 ± 3.3 | 34.3 ± 6.4 |

The effect of PNA in wt mice was analysed by an unpaired Student's t-test. wt PNA mice were compared to APNhet PNA to determine if there was an additive effect of adiponectin deficiency. Data are expressed as mean ± SEM.
